# A machine learning approach to define antimalarial drug action from heterogeneous cell-based screens

**DOI:** 10.1101/2019.12.19.882480

**Authors:** George W. Ashdown, Michelle Dimon, Minjie Fan, Fernando Sánchez-Román Terán, Katrin Witmer, David C. A. Gaboriau, Zan Armstrong, Jon Hazard, D. Michael Ando, Jake Baum

**Affiliations:** Department of Life Sciences, Imperial College London, London, UK; Facility for Imaging by Light Microscopy, Imperial College London, London, UK; Google Research, Applied Science Team

**Keywords:** Artificial intelligence, deep neural networks, high-throughput screening, *Plasmodium falciparum*, artemisinin-resistance, mitochondrial inhibitors, ATP4ase inhibitors

## Abstract

Drug resistance threatens the effective prevention and treatment of an ever-increasing range of human infections. This highlights an urgent need for new and improved drugs with novel mechanisms of action to avoid cross-resistance. Current cell-based drug screens are, however, restricted to binary live/dead readouts with no provision for mechanism of action prediction. Machine learning methods are increasingly being used to improve information extraction from imaging data. Such methods, however, work poorly with heterogeneous cellular phenotypes and generally require time-consuming human-led training. We have developed a semi-supervised machine learning approach, combining human- and machine-labelled training data from mixed human malaria parasite cultures. Designed for high-throughput and high-resolution screening, our semi-supervised approach is robust to natural parasite morphological heterogeneity and correctly orders parasite developmental stages. Our approach also reproducibly detects and clusters drug-induced morphological outliers by mechanism of action, demonstrating the potential power of machine learning for accelerating cell-based drug discovery.

**One Sentence Summary:** A machine learning approach to classifying normal and aberrant cell morphology from plate-based imaging of mixed malaria parasite cultures, facilitating clustering of drugs by mechanism of action.

---

Cell-based screens have significantly advanced our ability to find new drugs (*1*). However, most screens are unable to predict the mechanism of action (MoA) of identified hits, necessitating years of follow up post-discovery. In addition, even the most complex screens frequently discover hits against cellular processes that are already targeted (*2*). Limitations in finding new targets is becoming especially important in the face of rising antimicrobial resistance across bacterial and parasitic infections. This rise in resistance is driving increasing demand for screens that can intuitively find new antimicrobials with novel MoAs. Demand for innovation in drug discovery is exemplified in efforts targeting *Plasmodium falciparum*, the parasite that causes malaria. Malaria disease continues to be a leading cause of childhood mortality, killing nearly half a million children each year (*3*). Drug resistance has emerged to every major antimalarial to date including rapidly emerging resistance to frontline artemisinin-based combination therapies (*4*). Whilst there is a healthy pipeline of developmental antimalarials, many target common processes (*5*) and may therefore fail quickly because of prevalent cross-resistance. Thus, solutions are urgently sought for the rapid identification of new drugs that have a novel MoA at the time of discovery.

Parasite cell morphology within the human red blood cell (RBC) contains inherent MoA predictive capacity. Intracellular parasite morphology can distinguish broad stages along the developmental continuum of the asexual parasite (responsible for all disease pathology). This developmental continuum includes early development (early and late ring form), feeding (trophozoite), genome replication or cell division (schizont) and extra-cellular emergence (merozoite) (see (*6*) for definitions). As such, drugs targeting a particular stage should manifest a break in the continuum. Morphological variation in the parasite cell away from the continuum of typical development may also aid drug MoA prediction if higher information granularity can be generated during a cell-based screen. Innovations in automated fluorescence microscopy have markedly expanded available data content in cell imaging (*7*). By using multiple intracellular markers, an information-rich landscape can be generated from which morphology and potentially drug MoA can be deduced. This increased data content is, however, currently inaccessible both computationally and because it requires manual expert-level analysis of cell morphology. Thus, efforts to use cell-based screens to find drugs and define their MoA in a time efficient manner are still limited.

Machine learning (ML) methods offer a powerful alternative to manual image analysis, particularly deep neural networks (DNNs) that can learn to represent data succinctly. To date, supervised ML has been the most successful application for classifying imaging data, commonly based on binning inputs into discrete, human-defined outputs. Supervised methods using this approach have been applied to study mammalian cell morphologies (*8, 9*) and host-pathogen interactions (*10*). However, discrete outputs are poorly suited for capturing a continuum of morphological phenotypes, such as those that characterize either malaria parasite development or compound induced outliers, since it is difficult or impossible to generate labels of all relevant morphologies *a priori*. A cell imaging approach is therefore needed that can function with minimal discrete human-derived training data before computationally defining a continuous analytical space, which mirrors the heterogeneous nature of biological space.

Here we have created a semi-supervised model that discriminates diverse morphologies across the asexual lifecycle continuum of the malaria parasite *Plasmodium falciparum*. By receiving input from a Deep Metric Network (*11*) trained to represent similar consumer images as nearby points in a continuous coordinate space (an embedding). Our DNN can successfully define unperturbed parasite development with a much finer information granularity than human labelling alone. Importantly, the same DNN can quantify anti-malarial drug effects both in terms of lifecycle distribution changes (e.g. killing specific parasite stage(s) along the continuum) and morphological phenotypes or outliers not seen during normal asexual development. Combining lifecycle and morphology embeddings enabled the DNN to group compounds by their MoA without directly training the model on these morphological outliers. This DNN analysis approach towards cell morphology therefore addresses the combined needs of high-throughput cell-based drug discovery that can rapidly find new hits and predict MoA at the time of identification.

## Results

Using machine learning (ML), we set out to develop a high-throughput, cell-based drug screen that can define cell morphology and drug mechanism of action (MoA) from primary imaging data. From the outset we sought to embrace asynchronous (mixed stage) asexual cultures of the human malaria parasite, *Plasmodium falciparum*, devising a semi-supervised deep neural network (DNN) strategy that can analyse fluorescence microscopy images. The workflow is summarised in **Figure 1a-c**.

**Figure 1:**
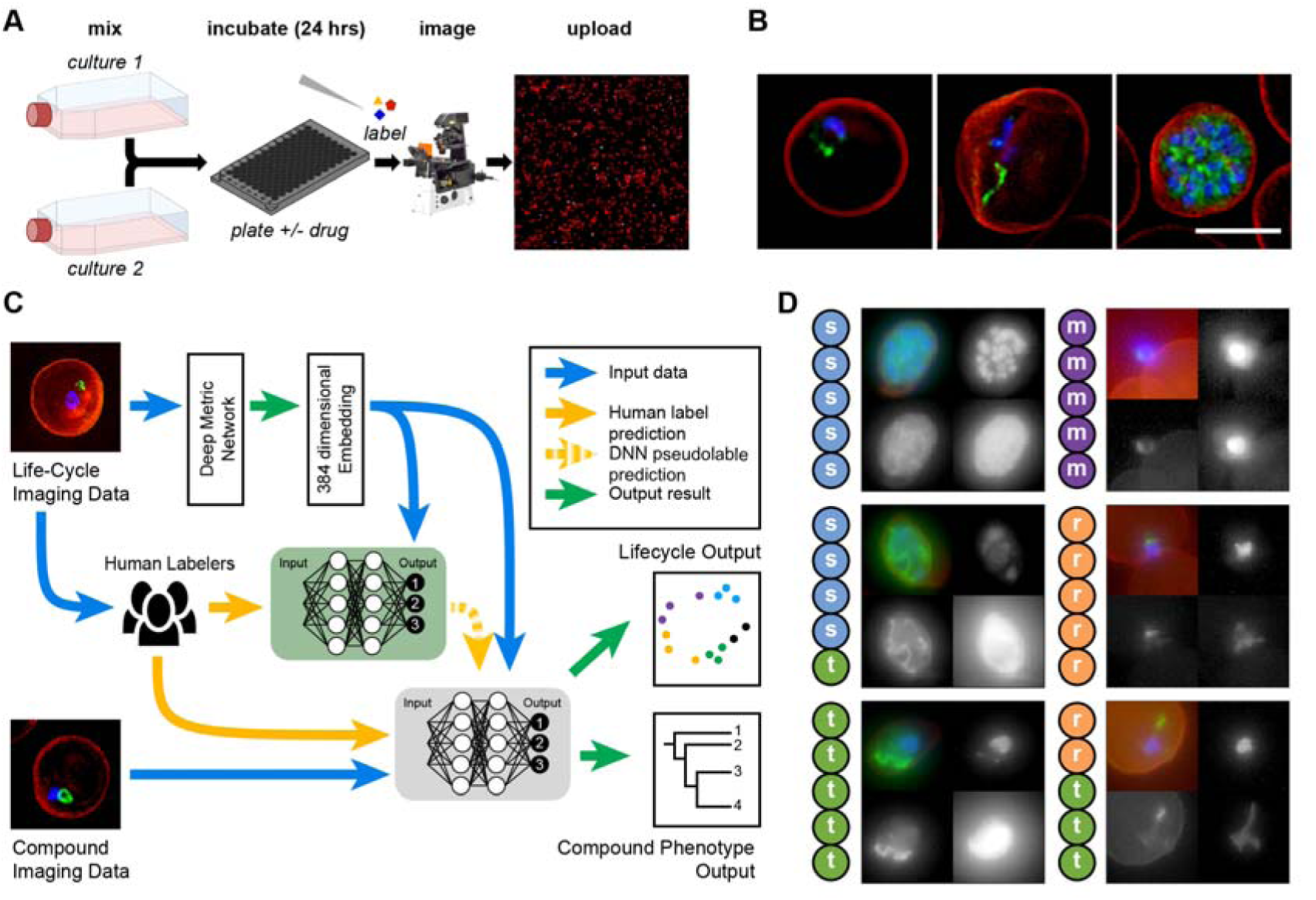
Experimental workflow. **A**) To ensure all lifecycle stages were present during imaging and analysis, two transgenic malaria cultures were combined (see Methods), these samples were incubated with or without drugs before being fixed and stained for automated multi-channel high-resolution, high-throughput imaging. Resulting datasets (**B**) contained parasite nuclei (blue), cytoplasm (not shown) and mitochondrion (green) information, as well as the red blood cell plasma membrane (red) and brightfield (not shown). These images were processed for machine learning analysis (**C**) with parasites segregated from full field of views using the nuclear stain channel, before transformation into embedding vectors. Two networks were used; the first (green) was trained on canonical examples from human labelled imaging data, providing machine learning derived labels (pseudolabels) to the second semi-supervised network (grey) which predicted lifecycle stage and compound phenotype. Example images from human labelled datasets (**D**) shows disagreement can occur between human labellers when categorising parasite stages (s = schizont, t = trophozoite, r = ring, m = merozoite). Scale bar = 5µm

### Overview of the approach: *Gathering human labels*

The *P. falciparum* lifecycle commences when free micron-sized parasites (called merozoites) target and invade human red blood cells (RBCs). During the first 8-12 hours post-invasion, the parasite is referred to as a “ring”, describing its diamond ring-like morphology (**Figure 1b, left**). The parasite then feeds extensively (trophozoite-stage, (**Figure 1b, middle**)), undergoing rounds of DNA replication and eventually divides into ∼20 daughter cells (the schizont-stage (**Figure 1b, right**)), which precedes merozoite release back into circulation (*6*). This discrete categorisation belies a continuum of morphologies between the different stages.

The morphological continuum of asexual development represents a challenge when teaching ML models, as definitions of each stage will vary between experts (**Figure 1d**). To embrace this, multiple human labels were collected. High resolution 3D images of transgenic 3D7 parasites were acquired using a widefield fluorescence microscope (**see Materials and Methods**), capturing brightfield, DNA (DAPI [4′,6-diamidino-2-phenylindole]), cytoplasm (constitutively expressed GFP), mitochondria (Mitotracker™) and the RBC membrane (fluorophore conjugated Wheat Germ Agglutinin, WGA). 3D datasets were converted to 2D images using maximum intensity projection. Brightfield was converted to 2D using both maximum and minimum projection, resulting in 6 channels of data for the ML. 5382 labels were collected from human experts, populating the categories of ring, trophozoite, schizont, merozoite, cluster-of- merozoites (multiple extracellular merozoites attached post RBC rupture), or debris. For initial validation and as a test of reproducibility between experts, an additional 448 parasites were collected, each labelled by 5 experts (**Figure 1d**).

### Overview of the approach: *Learning a data-driven representation of malaria phenotypes*

Image patches containing parasites within the RBC or after merozoite release were transformed into input embeddings using the Deep Metric Network architecture originally trained on consumer images (*11*) and previously shown for microscopy images (*12*). Embeddings are vectors of floating-point numbers representing a position in high-dimensional space, trained so related objects are located closer together. For our purposes, each image channel was individually transformed into an embedding of 64 dimensions before being concatenated to yield one embedding of 384 dimensions per parasite image.

Embeddings generated from parasite images were next transformed using a two-stage workflow to represent either *On-cycle* (for mapping the parasite lifecycle continuum) or *Off-cycle* effects (for mapping morphology or drug induced outliers). Initially, an ensemble of fully connected two-layer DNN models was trained on the input embeddings to predict the categorical human lifecycle labels for DMSO controls. This training gave the DNN robustness to control for sample heterogeneity and, as such, sensitivity for identifying unexpected results (outliers). The ensemble was built from three pairs of fully supervised training conditions (six total models). Each network pair was trained on separate (non-overlapping) parts of the training data, providing an unbiased estimate of the model prediction variance.

After initial training, the supervised DNN predicted its own labels (i.e. pseudolabels) for previously unlabelled examples. As with human derived labels, DNN pseudolabelling was restricted to DMSO controls (with high confidence) to preserve the model’s sensitivity to *Off-cycle* outliers (which would not properly fit into *On-cycle* outputs). High confidence was defined as images given the same label prediction from all six models and when all models were confident of their own prediction (defined as twice the random probability).

A new ensemble of models was then trained using the combination of human-derived labels and DNN pseudolabels. The predictions from this new ensemble were averaged to create the semi-supervised model.

### Validation of the *Plasmodium falciparum* lifecycle continuum

The semi-supervised model was first used to represent the normal (*On-cycle*) lifecycle continuum. We selected the subset of dimensions in the unnormalized final prediction layer that corresponded to merozoites, rings, trophozoites, and schizonts. This was projected into 2D space using principal component analysis (PCA) and shifted such that its centroid was at the origin. This resulted in a continuous variable where angles represent lifecycle stage progression, referred to as Angle-PCA. This Angle-PCA approach permitted the full lifecycle to be observed as a continuum with example images (**Figure 2a**) and 2D projection (**Figure 2b**) following the expected developmental order of parasite stages. Importantly, this ordered continuum manifested itself without specific constraints being imposed, except those provided by the categorical labels from human experts (see **Supplemental Note 1**).

**Figure 2:**
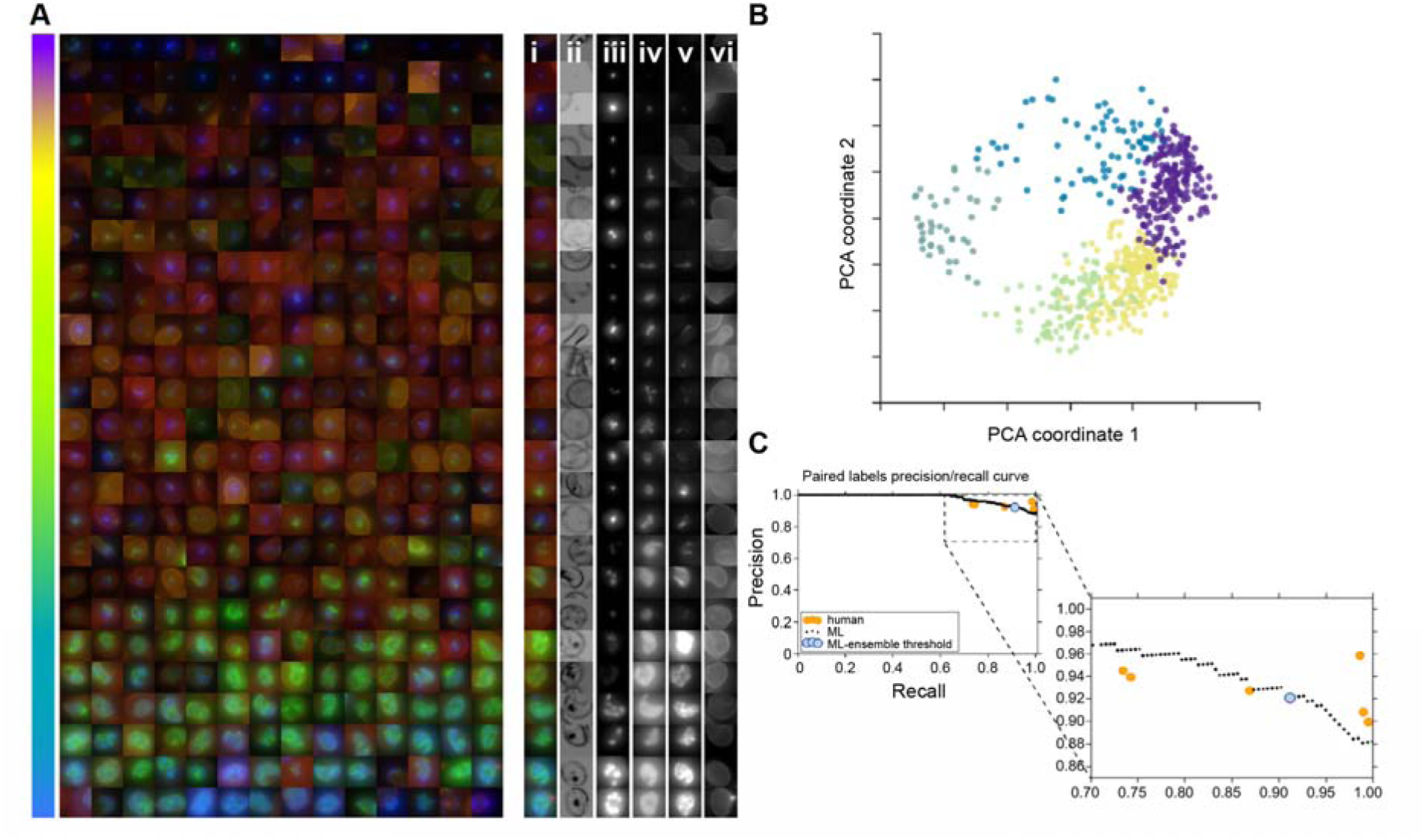
Machine learning continuum of parasite stages. After learning from canonical human-labelled examples and filtering debris and other outliers, the remaining lifecycle data from asynchronous cultures was successfully ordered by the model. The parasites shown are randomly selected DMSO control parasites from multiple imaging runs, sorted by Angle-PCA (**A**). The coloured, merged images show red blood cell membrane (red), mitochondria (green) and nucleus (blue). For a subset of parasites on the right, the coloured merged image plus individual channels are shown: (i) merged, (ii) brightfield min projection, (iii) nucleus, (iv) cytoplasm, (v) mitochondria, (vi) red blood cell membrane (brightfield max projection was also used in ML but not shown here). The model sorts the parasites in lifecycle stage order, despite heterogeneity of signal due to nuances such as imaging differences between batches. The order of the parasites within the continuum seen in (A) is calculated from the angle within the circle created by projecting model outputs using principal component analysis, creating a 2D scatterplot (**B**). This represents a progression through the life-cycle stages of the parasite, from merozoite (purple) through rings (yellow), trophozoites (green), schizonts (dark green) and finishing with cluster of merozoites (blue). The precision-recall curve (**C**) shows that human labellers and the model have equivalent accuracy in determining the earlier/later parasite in pairs. The consensus of the human labellers was taken as ground truth, with individual labellers (orange) agreeing with the consensus on 89.5-95.8% of their answers. Sweeping through the range of ‘too close to call’ values with the ML model yields the ML curve shown in black. Setting this threshold to 0.11 radians, the median angle variance across the individual models used in the ensemble, yields the blue dot.

To validate the accuracy of the continuous lifecycle prediction, pairs of images were shown to human labellers to define their developmental order (earlier/later) with the earliest definition being the merozoite stage. Image pairs assessed also included those considered indistinguishable (i.e. ‘too close to call’). Of the 295 pairs selected for labelling, 276 measured every possible pairing between 24 parasites while the remaining 19 pairs were specifically selected to cross the trophozoite/schizont boundary. Human expert agreement with the majority consensus was between 89.5% and 95.8%, with parasite pairs called equal (too close to call) 25.7% to 44.4% of the time. These paired human labels had more consensus than the categorical (merozoite, ring, trophozoite, schizont) labels which had between 60.9-78.4% agreement between individual human labels and the majority consensus.

The Angle-PCA projection results provide an ordering along the lifecycle continuum, allowing us to compare this sort-order to that by human experts. With our ensemble of six models, we could also evaluate the consensus and variation between angle predictions for each example. The consensus between models for relative angle between two examples was greater than 96.6% (and an area under the precision-recall curve score of 0.989), and the median angle variation across all labelled examples was 0.11 radians. The sensitivity of this measurement can be tuned by selecting a threshold for when two parasites are considered equal, resulting in a precision-recall curve (**Figure 2c**). When we use the median angle variation of the model as the threshold for examples that are too close to call, we get performance (light blue point) that is representative of the human expert average.

These results demonstrate that our semi-supervised model successfully identified and segregated asynchronous parasites and infected RBC’s from images which contain >90% uninfected RBCs (i.e. < 10% parasitaemia) and classifies parasite development logically along the *P. falciparum* asexual lifecycle.

### Quantifying *On-cycle* drug effects

Having demonstrated the semi-supervised model can classify asynchronous lifecycle progression consistently with fine-granularity, the model was next applied to quantify *on-cycle* differences (i.e. lifecycle stage-specific drug effects) in asynchronous, asexual cultures treated with known anti-malarial drugs. Two drug treatments were initially chosen that give rise to aberrant cellular development: the ATP4ase inhibitor KAE609 (also called Cipargamin) (*13*); and the mitochondrial inhibiting combinational therapy of atovaquone and proguanil (*14*) (herein referred to as AP). KAE609 reportedly induces cell swelling (*15*) while AP reduces mitochondrial membrane potential (*16*). Drug treatments were first tested at standard screening concentrations (2µM) for two incubation periods (6 hrs and 24 hrs). Next drug dilutions were carried out to test the semi-supervised model’s sensitivity to lower concentrations, using IC50’s of each compound (**Supp. Table 1**). IC50 and 2µM datasets were processed through the semi-supervised model and overlaid onto DMSO control data as a histogram, to explore *on-cycle* drug effects (**Figure 3**). KAE609 treatment exhibited a consistent skew towards ring stage parasite development (8-12 hrs post-RBC-invasion, **Figure 3**) without an increase within this stage of development, while the AP treatment led to reduced trophozoite stages (∼12-30 hrs post-RBC-invasion **Figure 3**). This demonstrates the fine-grained continuum has the sensitivity to detect whether drugs affect specific stages of the parasite lifecycle.

**Figure 3:**
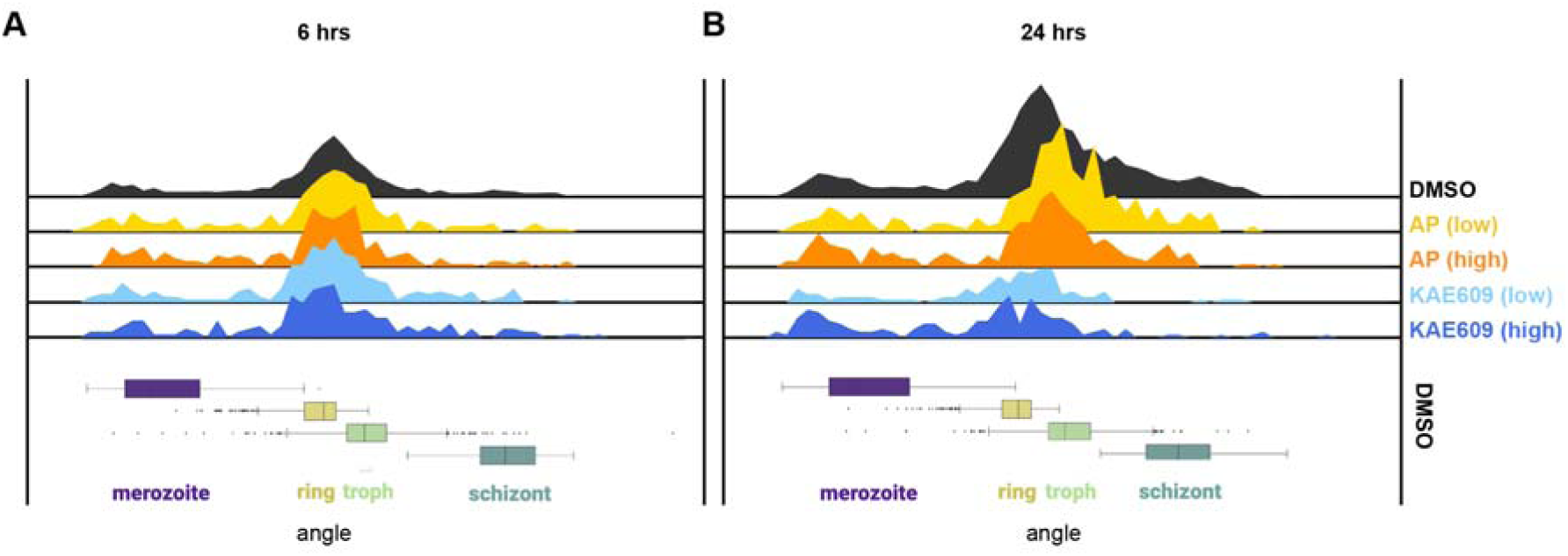
Quantifying *on-cycle* drug effects. Asynchronous *Plasmodium falciparum* cultures were treated with the ATPase4 inhibitor KAE609 or the combinational mitochondrial treatment of atovaquone and proguanil (AP) with samples fixed and imaged 6 hr (**A**) and 24 hrs (**B**) after drug additions. Top panels show histograms indicating the number of parasites across lifecycle continuum. Compared to DMSO controls (topmost black histogram) both treatments demonstrated reduced parasite numbers after 24 hr. Shown are four drug/concentration treatment conditions: low dose AP (yellow), high dose AP (orange), low dose KAE609 (light blue), high dose KAE609 (dark blue). Box plots below demonstrate life-cycle classifications in the DMSO condition of images from merozoite (purple) through rings (yellow), trophozoites (green) and finishing with schizonts (dark green).

### Identifying *Off-cycle* drug effects on parasite morphology

The improved information granularity was extended to test if the model could identify drug-based morphological phenotypes (*off-cycle*), towards determination of MoA. Selecting the penultimate 32-dimensional layer of the semi-supervised model meant, unlike the Angle-PCA model, outputs were not restricted to discrete *on-cycle* labels, but instead represented both *on-* and *off-cycle* changes. This 32-dimensional representation is referred to as the morphology embedding.

Parasites were treated with one of 11 different compounds, targeting either PfATP4ase (ATP4) or mitochondria (MITO) plus DMSO controls (**Supp. Table 1**). The semi-supervised model was used to evaluate three conditions: random; where compound labels were shuffled, Angle-PCA; where the two PCA coordinates are used, and full embedding; where the 32-dimensional embedding was combined with the Angle-PCA. To add statistical support that enables compound level evaluation, a bootstrapping of the analysis was performed, sampling a subpopulation of parasites 100 times.

As expected, the randomized labels led to low accuracy (**Figure 4a**), serving as a baseline for the log odds (probability). When using the 2-dimensional Angle-PCA (*on-cycle*) information, there was a significant increase over random in the log odds ratio (**Figure 4a**). This represents the upper-bound information limit for binary live/dead assays due to their insensitivity to parasite stages. When using the combined full embedding, there was a significant log odds ratio increase over both the random and Angle-PCA conditions (**Figure 4a**). To validate that this improvement was not a consequence of having a larger dimensional space compared to the Angle-PCA, an equivalent embedding from the fully supervised model trained only on expert labels (and not on pseudolabels) demonstrated approximately the same accuracy and log odds ratio as Angle-PCA. Thus, our semi-supervised model can create an embedding sensitive to the phenotypic changes under distinct MoA compound treatment.

**Figure 4:**
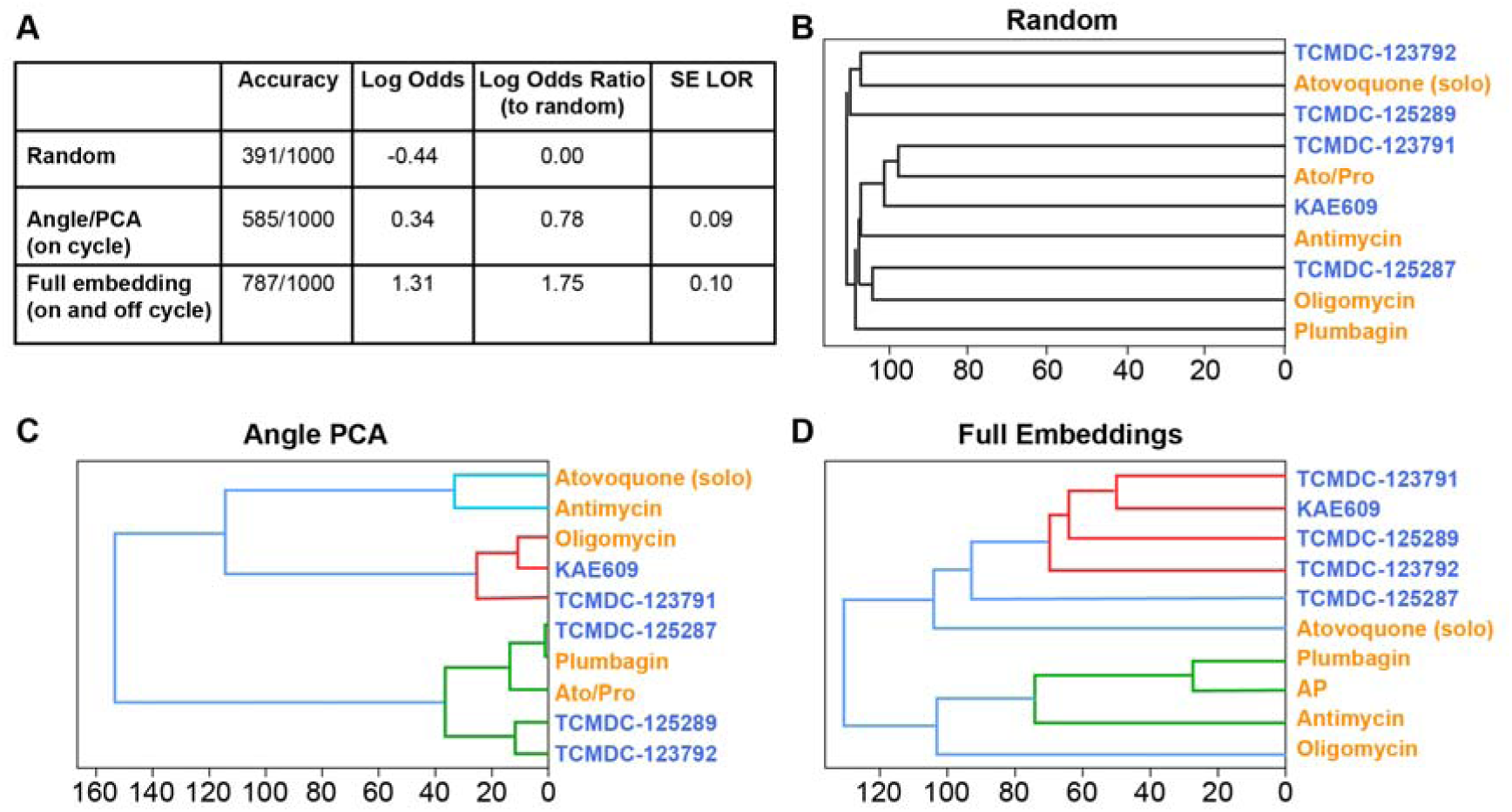
Quantifying *off-cycle* drug effects. To better define drug effect on *Plasmodium falciparum* cultures, 5 mitochondrial (red text) and 5 PfATP4ase (blue text) compounds were used, after a 24 hr incubation images were collected and analysed by the semi-supervised model. To test performance various conditions were used (**A**). For random, images and drug names were scrambled, leading to the model incorrectly grouping compounds based on known mechanism of action (**B**). Using life-cycle stage definition (as with **Figure 3**) the model generated improved grouping of compounds (**C**) versus random. Finally, by combining the life-cycle stage information with the penultimate layer (morphological information, prior to life-cycle stage definition) of the model led to correct segregation of drugs based on their known mechanism of action (**D**).

To better understand drug MoA, we evaluated how the various compounds were grouped together by the three approaches (random, Angle-PCA, and morphology embedding), performing a hierarchical linkage dendrogram (**Figure 4b-d**). The random approach shows, as expected, the different compounds do not reveal MoA similarities. For the Angle-PCA output, the MITO inhibitors Atovaquone and Antimycin are grouped similarly, but the rest of the clusters are a mixture of compounds from the two MoA groups. Finally, the morphology embedding gave rise to an accurate separation between the two groups of compounds having different MoA. One exception for grouping was Atovaquone (when used alone), which was found to poorly cluster with either group (branching at the base of the dendrogram) (**Figure 4d**). This result is likely explained by the drug dosages used, as Atovaquone is known to have a much-enhanced potency when used in combination with Proguanil (*16*).

The semi-supervised model was able to consistently cluster MITO inhibitors away from ATP4ase compounds in a dimensionality that suggested a common MoA. Our semi-supervised model can therefore successfully define drug efficacy *in vitro* and simultaneously assign a potential drug MoA from asynchronous (and heterogeneous) *P. falciparum* parasite cultures, using an imaging-based screening assay with high-throughput capacity.

## Discussion

Driven by the need to accelerate novel antimalarial drug discovery with defined mechanism of action (MoA) from phenotypic screens, we applied machine learning (ML) to images of asynchronous *P. falciparum* cultures. This semi-supervised ensemble model could identify effective drugs and cluster them according to MoA, based on lifecycle stage (*on-cycle*) and morphological outliers (*off cycle*).

Recent image-based ML approaches have been applied to malaria cultures but have, however, focussed on automated diagnosis of gross parasite morphologies from either Giemsa or Leishman-stained samples (*17-19*), rather than phenotypic screening for drug MoA. ML of fluorescence microscopy images have reported malaria identification of patient-derived blood smears (*20*) and the use of nuclear and mitochondrial specific dyes for stage categorization and viability (*21*), though without an AI approach. Previous unsupervised and semi-supervised ML approaches have been applied to identify phenotypic similarities in other biological systems, such as cancer cells (*12, 22-24*), but none have addressed the challenge of capturing the continuum of biology within the heterogeneity of control conditions. We therefore believe our study represents the first time that high-resolution imaging data has been used beyond diagnostics to predict the lifecycle continuum of a cell type (coping with biological heterogeneity), as well as using this information to indicate drug-induced outliers and successfully group these towards drug MoA.

Through semi-supervised learning, only a small number of human-derived discrete but noisy labels from asynchronous control cultures were required for our DNN method to learn and distribute data as a continuous variable, with images following the correct developmental order. By reducing expert human input, which can lead to image-identification bias (see **Supplemental Note 1**), this approach can control for inter-expert disagreement and is more time efficient. This semi-supervised DNN therefore extends the classification parameters beyond human-based outputs, leading to finer information granularity learned from the data automatically through pseudolabels. This improved information, derived from high resolution microscopy data, permits the inclusion of subtle but important features to distinguish parasite stages and phenotypes that would otherwise be unavailable.

Our single model approach was trained on lifecycle stages through embedding vectors, whose distribution allows identification of two readouts, *on-cycle* (sensitive to treatments which slow the lifecycle or kill a specific parasite stage) and *off-cycle* (sensitive to treatments which cluster away from control distributions). We show that this approach with embeddings was sensitive to stage specific effects at IC50 drug concentrations (**Figure 3**), much lower than standard screening assays. Drug based outliers were grouped in a MoA-dependent manner (**Figure 4**), with data from similar compounds grouped closer than data with unrelated mechanisms.

The simplicity of fluorescence imaging means this method could be applied to different sub-cellular parasite features, potentially improving discrimination of cultures treated with other compounds. Additionally, imaging the sexual (gametocyte) parasite stages with and without compound treatments will build on the increasing need for drugs which target multiple stages of the parasite lifecycle (*25*). Current efforts to find drugs targeting the sexual stages of development are hampered by the challenges of defining MoA from a non-replicating parasite lifecycle stage (*25*). This demonstrates the potential power of a mechanism of action approach, applied from the outset of their discovery, based simply on cell morphology.

In the future, we envisage *on-cycle* effects could elucidate the power of combinational treatments (distinguishing treatments targeting different lifecycle stages) for a more complete therapy. Using *off-cycle* this approach could identify novel combinational treatments based on MoA. Due to the sample preparation simplicity, this approach is also compatible with using drug resistant parasite lines.

New drugs against malaria are seen as a key component of innovation required to ‘bend-the-curve’ towards the disease’s eradication or risk a return to pre-millennium rates (*3, 26*). Seen in this light, application of ML-driven screens should enable the rapid, large scale screening and identification of drugs with concurrent determination of predicted MoA. Since ML-identified drugs will start from the advanced stage of predicted MoA, these should bolster the much-needed development of new chemotherapeutics for the fight against malaria.

## Funding

This work was funded by a grant from the Bill & Melinda Gates Foundation (OPP1181199) with additional support coming from Wellcome (Investigator Award to JB, 100993/Z/13/Z JB). We acknowledge the Facility for Imaging by Light Microscopy (FILM) at Imperial College London, which runs a Nikon High-Content microscope platform funded by Wellcome (104931/Z/14/Z). We would like to thank several colleagues for their help with expert labelling of data, including, Irene Garcia Barbazan, Tom Blake, Alisje Churchyard, Farah Dahalan, Mark Wilkinson, Katherine Wright and Sabrina Yahiya.

## Author contributions

GWA, MD, MA and JB designed the experiments, FSRT cultured the parasites for imaging, KW generated the 3D7/P230p-sfGFP parasite line, DG automated the imaging, GWA optimised and carried out sample preparation and imaging, MD, MF, MA and JH optimised and carried out the machine learning workflow and data analysis. All authors contributed to writing the manuscript.

## Competing interests

The authors declare no conflicts of interest or competing interests.

## Data and materials availability

Original imaging data, experimental metadata and pseudolabels will be available from the Image Data Repository (pending upload).

## Supplementary Materials

Materials and Methods

Supplementary Note 1

Figure S1

Tables S1

