## Supplementary Materials for "A machine learning approach to define antimalarial drug action from heterogeneous cell-based screens"

**This PDF file includes:**

Materials and Methods

Supplementary Text

Fig. S1

Table S1

### Materials and Methods

#### *Generation of GFP+ 3D7 parasite line*

To generate parasite line 3D7/P230p-sfGFP, 3D7 ring stages were transfected with both plasmids pkiwi003 (p230p-bsfGFP) and pDC2-cam-co.Cas9-U6.2-hDHFR _P230p (50ug each, **Supplementary Figure S1**) following standard procedures (*27*) and selected on 4nM WR99210 (WR) for 10 days. pDC2-cam-co.Cas9-U6.2-hDHFR _P230p encodes for Cas9 and the guide RNA for the P230p locus. pkiwi003 comprises the repair sequence to integrate into the P230p locus after successful double strand break induced by the Cas9. pkiwi003 (p230p-bsfGFP) was obtained by inserting two PCR fragments both encoding parts of P230p (PF3D7_0208900) consecutively into the pBluescript SK(-) vector with XhoI/HindIII and NotI/SacI, respectively. sfGFP together with the hsp70 (bip) 5' UTR was PCR amplified from pkiwi002 and cloned into pkiwi003 with HindIII/NotI. pkiwi002 is based on pBSp230pDiCre (*28*), where the FRB and Cre60 cassette (including promoter and terminator) was removed with AfeI/SpeI, and the following linkers inserted: L1_F cctttttgcccccagcgctatataactagtACAAAAAGTATCAAG and L1_R CTTGATACTTTTTTGTactagttatatagcgctgggggcaaaaagg. In a second step, FKBP and Cre59 were removed with NheI/PstI and replaced by sfGFP, which was PCR amplified from pCK301 (*29*). pDC2-cam-co.Cas9-U6.2-hDHFR _P230p was obtained by inserting the guide RNA (AGGCTGATGAAGACATCGGG) into pDC2-cam-co.Cas9-U6.2-hDHFR (*30*) with BbsI. Integration of pkiwi003 into the P230p locus was confirmed by PCR using primers #99 (ACCATCAACATTATCGTCAG); #98 (TCTTCATCAGCCTGGTAAC) and #56 (CATTTACACATAAATGTCACAC) (**Supplementary Figure S1**).

#### *Plasmodium falciparum cell culture*

*P. falciparum* asexual parasites (3D7) were cultured at 37 °C [with a gas mixture of 90% N2, 5% O2 and 5% CO2] in human O+ erythrocytes under standard conditions (*31*), with RMPI-HEPES medium supplemented with 0.5% AlbuMAX-II. Two independent stocks (culture 1 and culture 2 [**Figure 1a**]) of 3D7 parasites were maintained in culture and synchronized separately with 5% D-Sorbitol on consecutive days to ensure acquisition of all stages of the asexual cycle on the day of sample preparation. Samples used for imaging derived from cultures harbouring an approximate 1:1:1 ratio of rings, trophozoites and schizonts, with a parasitaemia around 10%.

***Sample preparation and imaging***

Asexual cultures were diluted 50:50 in fresh media before 50nM MitoTracker™ CMXRos (ThermoFisher) was added for 20 mins at 37°C. Samples were then fixed in PBS containing 4% formaldehyde and 0.25% glutaraldehyde and placed on a roller at room temperature, protected from light for 20 mins. The sample was then washed 3× in PBS before 10 nM DAPI (4′,6-diamidino-2-phenylindole) and 5 µg/ml wheat germ agglutinin (WGA)- conjugated to AlexaFluor633 was added for 10 mins and protected from light. The sample was then washed 1x in PBS and diluted 1 in 30 in PBS before pipetting 100µL into each well of a CellVis (Mountain View, CA) 96-well plate.

Samples were imaged using a Nikon Ti-Eclipse widefield microscope and Hamamatsu EMCCD camera, with a 100× Plan Apo 1.4NA oil objective lens (Nikon), the NIS-Elements JOBS software package (Nikon) was used to automate the plate-based imaging. The five channels (brightfield, DNA (DAPI), cytoplasm (sfGFP-labeled), mitochondria (MitoTracker™), and RBC (WGA-633)) were collected serially at Nyquist sampling as a 6µm z-stack, with fluorescent excitation from the CoolLED light source. To collect enough parasite numbers per treatment, 32 fields of view (sites) were randomly generated and collected within each well, with treatments run in technical triplicate. Data was saved directly onto an external hard-drive for short-term storage and processing (see below).

***Image analysis***

The 3D images were processed via a custom macro using ImageJ and transformed into 2D maximum intensity projection images. Brightfield channels were also projected using the minimum intensity projection as this was found to improve analysis of the food vacuole and anomalies including double infections. Converting each whole-site image to per-parasite embedding vectors was performed as previously described (*12*), with some modifications: the Otsu threshold was set to the minimum of the calculated threshold or 1.25x the foreground mean of the image, and centres closer than 100 pixels were pruned. Each channel image was separately fed as a grayscale image into the Deep Metric Network for conversion into a 64-dimension embedding vector. The 6 embedding vectors (one from each fluorescent channel, and both min and max projections of the brightfield channel) were concatenated to yield a final 384 dimension embedding vector.

***Ground truth***

All labels were collected using the annotation tool originally built for collecting diabetic retinopathy labels (*32*).

For each set of labels gathered, tiled images were stitched together to create a collage for all parasites to be labelled. These collages contained both stains in grayscale as well as colour overlays to aid identification. Collages and a set of associated questions were uploaded to the annotation tool and human experts (Imperial College London) provided labels (answers). In cases where multiple experts labelled the same image, a majority vote was used to determine the final label.

Initial labels for training classified parasites into one of eleven classes: merozoite, ring, trophozoite, schizont, cluster of merozoites, multiple infection, bad image, bad patch (region of interest) location, parasite debris, unknown parasite inside an RBC, or other. Subsequent labels were collected with parasite debris classified further into: small debris remnant, cluster of debris, and death inside a red blood cell. For training, the following labels were dropped: bad image, bad patch location, unknown parasite inside an RBC, unspecified parasite debris, and other. For these labels, five parasites were randomly sampled from each well of experiments.

To validate the model performance, an additional 448 parasites were labelled by 5 experts. The parasites were selected from eight separate experimental plates, using only control image data (DMSO only).

Finally, paired labels were collected to validate the sort-order results. For these labels, the collage included two parasites and experts identified which parasite was earlier in the lifecycle, or whether the parasites were too close to call. Here, data from the 448 parasite validation set were used, limited to cases where all experts agreed the images were of a parasite inside a red blood cell. From this set, 24 parasites were selected and all possible pairings of these 24 parasites were uploaded as questions (24 choose 2 = 276 questions uploaded). In addition, another 19 pairs were selected that were near the trophozoite/schizont boundary, to enable angle resolution analysis.

***Data analysis***

To prepare the data for analysis, the patch embeddings were first joined with the ground truth labels for patches with labels.

Six separate models were trained on embeddings to classify asexual life cycle stages and normal anomalies such as multiple infection, cell death, and cellular debris. Each model was a two-layered (64 and 32 dimensions), fully connected (with ReLu non-linearities) neural network. To create training data for each of the six models, human-labelled examples were partitioned so that each example within a class is randomly assigned to one of four partitions. Each model was then trained on one of the six ways to select a pair from the four partitions. Training was carried out with a batch size of 128 for 1,000 steps using the Adam optimizer (*33*) with a learning rate of 2e-4. Following the initial training, labels were predicted on all unlabelled data using all six models, and for each class 400 examples were selected with the highest mean probability (and at least a mean probability of 0.4) and with a standard deviation of the probability less than 0.07 (which encompasses the majority of the predictions with labels). The training procedure was repeated with the original human-labels and predicted (pseudo-) labels to generate our final model. The logits are extracted from the trained model and a subspace representing the normal life cycle stages is projected using two-dimensional by PCA. The life cycle angle is computed as arctan(y/x), where x and y are the first and second coordinates of the projection, respectively.

For each drug with a certain dose and application duration, the evaluation of its effect is based on the histogram of the classified asexual life cycle stages, and finer binned stages obtained from the estimated life cycle angle.

### Supplementary Text

#### Supplementary Note 1: Human experts provide noisy initial labels to train supervised models

With typical mammalian cells, almost every normal cell should look approximately the same. Dividing cells look different but are sparse and generally dropped from analysis. *P. falciparum* shows dramatic morphological changes throughout its lifecycle. In order to successfully apply ML, especially when investigating drug induced changes, 'normal' needed to be defined throughout the lifecycle. One solution would be to create synchronized cultures at each stage and use these as the ground truth. In reality, even these synchronized cultures show parasite-to-parasite variability (a 'trophozoite-heavy' culture is still a mix of some late rings, some trophozoites, some early schizonts). Instead, asynchronous cultures were used and collected ground truth labels from human experts. This presents a segregation challenge for the ML when channel intensities range greatly (e.g. DAPI brightness between ring and schizont stages [Figure 1c]). Human labels enable training of a standard supervised random forest model to bin parasites into ring / trophozoite / schizont stages. However, these include increased levels of noise, especially away from canonical images, for example experts disagree about whether a parasite is a late ring or early trophozoite. A random forest trained on these labels also has disagreement with the held-out test dataset. It is unclear if this disagreement is because the human labels are noisy or because the random forest is poorly trained.

### Supplementary Figure S1: Generation of *P. falciparum* constitutively expressing sfGFP

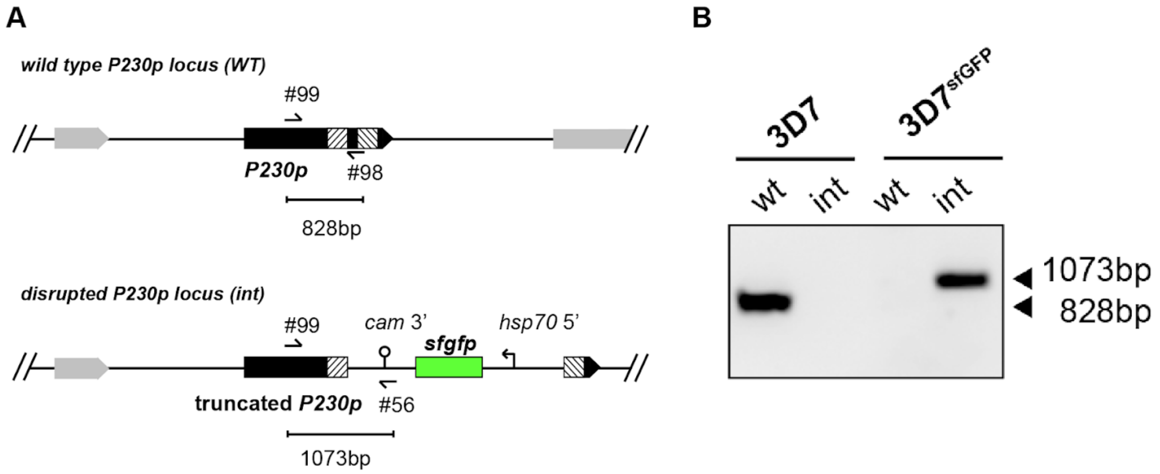


**A.** Schematic map of the *P230p* locus before (WT, top) and after integration of plasmid pkiwi003 (int, bottom). **B.** Integration PCR confirming integration of plasmid pkiwi003 into the P230p locus using primer pairs (#99/98 WT; #99/56 int) from (A).

**Supplementary Table S1. List of drugs used for screening phase.**

| Drug | Target / Effect | IC50 (nM) | Reference |
|---|---|---|---|
| Atovaquone + Proguanil hydrochloride | Mitochondria Electron Transport chain $bc_1$ Complex III inhibitor | 0.23 + 45 (in combination) | (*16*) |
| Oligomycin A | Mitochondrial ATP synthase Complex V inhibitor | 110 | PubChem (*34*) |
| Atovaquone | Mitochondria Electron Transport chain $bc_1$ Complex III (Q0 site) inhibitor | 0.27 | (*35*) and PubChem (*34*) |
| Antimycin A | Mitochondria Electron Transport chain $bc_1$ Complex III (Q1 site) inhibitor | 614 | (*36*) |
| Plumbagin | Succinate dehydrogenase (SDH) complex II inhibitor | 580 | (*37*) |
| KAE609 (cipargamin) | Inhibits ATP4ase, disrupting Na+ regulation in parasites | 1 | (*15*) |
| TCMDC-125287 | PfATP4ase inhibitor | 1700 | GSK report on PubChem (*34*) |
| TCMDC-125289 | PfATP4ase inhibitor | 50 | GSK report on PubChem (*34*) |
| TCMDC-123791 | PfATP4ase inhibitor | 970 | GSK report on PubChem (*34*) |
| TCMDC-123792 | PfATP4ase inhibitor | 200 | GSK report on Pubchem (*34*) |
